# A Kernel-Based Change Detection Method to Map Shifts in Phytoplankton Communities Measured by Flow Cytometry

**DOI:** 10.1101/2020.12.01.405126

**Authors:** Corinne Jones, Sophie Clayton, François Ribalet, E. Virginia Armbrust, Zaid Harchaoui

## Abstract

1. Automated, ship-board flow cytometers provide high-resolution maps of phytoplankton composition over large swaths of the world’s oceans. They therefore pave the way for understanding how environmental conditions shape community structure. Identification of community changes along a cruise transect commonly segments the data into distinct regions. However, existing segmentation methods are generally not applicable to flow cytometry data, as this data is recorded as “point cloud” data, with hundreds or thousands of particles measured during each time interval. Moreover, nonparametric segmentation methods that do not rely on prior knowledge of the number of species, are desirable to map community shifts.
2. We present CytoSegmenter, a kernel-based change-point estimation method for segmenting point cloud data that does not rely on parametric assumptions on the data distributions. Our method relies on a Hilbertian embedding of point clouds that allows us to work with point cloud data similarly to vectorial data. The change-point locations can be found using an efficient dynamic programming algorithm. The method can be used to automatically segment long series of underway flow cytometry data.
3. Through an analysis of 12 cruises, we demonstrate that CytoSegmenter allows us to locate abrupt changes in phytoplankton community structure. We show that the changes in community structure generally coincide with changes in the temperature and salinity of the ocean. We also illustrate how the main parameter of CytoSegmenter can be easily calibrated using limited auxiliary annotated data.
4. CytoSegmenter is publicly available and implemented in the programming language Python. The method is generally applicable for segmenting series of point cloud data from any domain. Moreover, it readily scales to thousands of point clouds, each containing thousands of points. In the context of underway flow cytometry data, it does not require prior clustering of particles to define taxa labels, eliminating a potential source of error. This represents an important advance in automating the analysis of large datasets now emerging in biological oceanography and other fields. It also allows for the approach to potentially be applied during research cruises.

## 1 Introduction

Determining the number and locations of abrupt changes in distribution of a sequence of observations has played an important role in analyzing ecological data. The study of change points in marine ecology and oceanography dates back several decades. Legendre et al. (1985) performed change-point detection on time series of zooplankton counts near the Mediterranean Sea and in a reservoir in Quebec in the late 1960s and 1970s, respectively. Since then, change points in time have been studied at a wide range of spatial and temporal scales. Spatially, this includes large-scale changes in environmental and/or biological variables measured in the North Pacific and North Atlantic (Mantua et al., 1997, Hare and Mantua, 2000, Friedland et al., 2016) and smaller-scale changes in the abundance of species in seas, lakes and estuaries (Gal and Anderson, 2010, Thomson et al., 2010, Alvarez-Fernandez et al., 2012, Weijerman et al., 2005). The time scale between change points depends on the causes of the changes. For example, studies have examined interdecadal changes the North Pacific related to changes in the climate, month-long changes caused by eutrophication (Pace et al., 2017), and daily changes due to vertical migration (Bianchi and Mislan, 2016).

While all of the aforementioned literature focuses on change points in time, it is equally possible to detect change points in space. For example, Nieuwhof et al. (2018) detected changes in water storage capacity as the distance from shellfish reefs increased, and Li et al. (2019) identified the boundary of the Antarctic Intermediate Waterway. In this work we draw on ideas from the nonparametric statistics and the machine learning literature and develop a retrospective change-point estimation method for series of point cloud data that can be applied to segment flow cytometry data.

Over the past three decades, flow cytometry has become instrumental in determining the distribution of phytoplankton communities (Sosik et al., 2010). Flow cytometry measures scattered light and fluorescence emissions of individual cells at rates of up to thousands of cells per second. Light scattering is proportional to cell size, and fluorescence is unique to the emission spectra of pigments; together, these parameters can be used to identify populations of phytoplankton with similar optical properties. Automated flow cytometers such as CytoBuoy (Dubelaar et al., 1999), FlowCytobot (Olson et al., 2003), and SeaFlow (Swalwell et al., 2011) have provided unprecedented views of the dynamics of phytoplankton across large temporal and spatial scales. The recent release of SeaFlow data collected underway during cruises conducted in the North Pacific Ocean offers unique opportunities to study how phytoplankton communities vary over space and time (Ribalet et al., 2019). Understanding how phytoplankton communities vary in time and space across ocean basins is critical for predicting how marine ecosystems will respond to future climate change.

Performing change-point analysis on underway flow cytometry data would allow us to segment a cruise into statistically distinct regions based on measures such as the scatter and fluorescence of individual particles. However, the underway flow cytometry data produced by instruments such as SeaFlow presents novel challenges not addressed by existing segmentation methods. The individual phytoplankton measurements are recorded with a temporal resolution of 3 min (roughly a 1 km spatial resolution). Hence, each observation corresponding to one time point can be viewed as a point cloud of individual phytoplankton measurements. Moreover, as the datasets collected during any given research cruise might contain more than 50 million particle measurements, the method must scale.

Methods that have been used in ecology and oceanography for detecting change points generally fall into one of three categories. First, there are methods that repeatedly perform hypothesis tests at every location in the time series and set a testing threshold to yield an appropriate number of change points (Page, 1954, Quandt, 1958, Srivastava and Worsley, 1986, Clarke, 1993, Matteson and James, 2014). Second, there are methods that estimate the locations of a large number of potential change points and then prune them using a penalty term or hypothesis testing (Gordon and Birks, 1972). Finally, there are methods that fit a single model to the time series that allows for an unknown number of changes at unknown times (Hamilton, 1990, Goldfeld and Quandt, 1973, Fearnhead, 2006). However, nearly all of these approaches assume that the data is either real-valued or vectorial. Applying such methods by reducing each point cloud to its mean as done by Hyrkas et al. (2015) results in the loss of important information regarding the distribution of the data in each point cloud. Of the cited methods, only the dissimilarity-based approaches of Clarke (1993) and Gordon and Birks (1972) could potentially be applied to data consisting of sequences of point clouds. Even if these methods were used, an appropriate dissimilarity measure would still need to be chosen and a principled method for determining the number of changes would need to be proposed.

In this work we develop a nonparametric statistical method for identifying abrupt changes in large-scale series of point clouds of measurements. The method is nonparametric in that it does not require the modeler to specify a parametric family of probability distributions. It therefore has the advantage that, given enough samples, it can detect any change in probability distribution. Moreover, it is capable of scaling to sequences of thousands of point clouds, each with thousands of points. We demonstrate the approach by segmenting SeaFlow data along cruise tracks. We also assess how closely estimates of biological change points coincide with physical shifts to better understand the role of the physical ocean environment in controlling phytoplankton community structure and to estimate the number of biological change points.

## 2 Materials and methods

### 2.1 SeaFlow data

To date, the SeaFlow flow cytometer has been used on more than 60 cruises in the North Pacific Ocean. In this work we focus on 12 of those cruises. These cruises are all of the cruises that are neither coastal cruises nor cruises around the Hawaiian islands and for which data is available. The locations of the cruises are depicted in Fig. 1 and some characteristics of the cruises may be found in Table 1.

**Figure 1:**
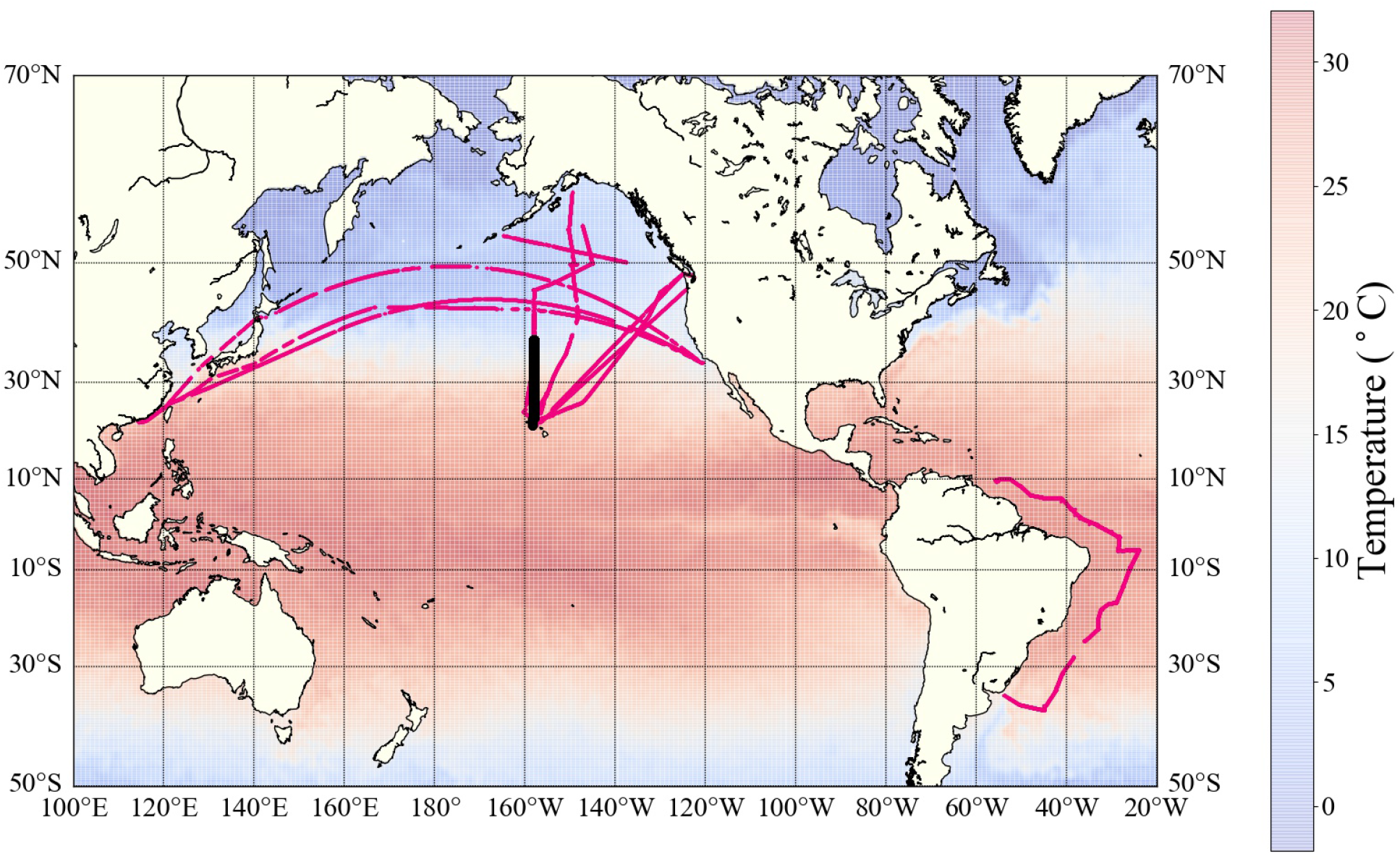
Locations of the cruises analyzed in this paper, overlaid on sea surface temperature data from April 26, 2016.^1^ In this work we primarily focus on KOK1606, the cruise in black in the middle of the map, which took place from April 20-May 4, 2016.

**Table 1:** Details of the cruises used. The column “# changes” provides the number of change points in the annotated physical data.

| Cruise | Location | Length (km) | # point clouds | # particles | # changes |
| --- | --- | --- | --- | --- | --- |
| KM1502 | Portland - Hawaii | 3,987 | 3,682 | 6.3 million | 44 |
| KM1712 | Hawaii - Alaska | 5,910 | 10,455 | 12.4 million | 48 |
| KM1713 | Alaska - Hawaii | 4,458 | 8,877 | 13.7 million | 27 |
| KN210-04 | South Atlantic | 17,436 | 18,260 | 56.7 million | 92 |
| KOK1606 | Subtropical front | 4,006 | 6,567 | 2.3 million | 60 |
| MGL1704 | Subtropical front | 4,939 | 7,054 | 5.8 million | 32 |
| TN248 | Gulf of Alaska | 2,049 | 1,362 | 0.2 million | 28 |
| TN271 | Seattle - Hawaii | 4,527 | 3,552 | 1.0 million | 41 |
| TN292 | Seattle - Hawaii | 3,905 | 3,120 | 2.8 million | 30 |
| Tokyo_1 | North Pacific | 11,384 | 4,871 | 0.9 million | 83 |
| Tokyo_2 | North Pacific | 11,485 | 5,159 | 1.5 million | 48 |
| Tokyo_3 | North Pacific | 11,664 | 5,627 | 3.1 million | 118 |

The data we use consists of two components. First, there is physical data, which consists of measurements of sea surface temperature and salinity collected at 3-minute intervals throughout the cruises from the ship’s underway thermosalinograph system. Second there is biological data, which consists of measurements of light scatter and fluorescence emissions of individual particles. The biological data is organized into point clouds recorded every 3 min, and each post-processed point cloud contains measurements of the cytometric characteristics of between 100 and 10,000 particles ranging from 0.5 to 5 microns in diameter. The volume of data in any given file depends on the total abundance of phytoplankton within the sampled region. Each particle is characterized by two measures of fluorescence emission (chlorophyll and phycoerythrin), its light scatter, and its label (identified based on a combination of manual gating and a semi-supervised clustering method) (Ribalet et al., 2019). Note that we use the particle labels only for verification of our approach.

The data is cleaned as follows. For the physical data we first remove the observation times that are not in chronological order. Next, we remove observations for which the temperature is unavailable or larger than 60°C. Furthermore, we remove outlier observations for which the salinity is less than 10 or larger than 60 PSU. Finally, we remove successive files that were recorded at times when the ship had not moved more than 0.1 km. For the biological data we remove files for which the physical data is not available. We also delete the entries corresponding to added calibration beads rather than phytoplankton.

One downside to this dataset is that there are no ground-truth change points in either the biological data or the physical data that could be used to evaluate our method. To ameliorate this problem, we manually annotated change points in the physical data using the following reasoning. Changes in phytoplankton community structure often occur when different water parcels mix together. A mixing event of different water masses can be identified by studying changes in water temperature and salinity. The mass of a water parcel (defined by its temperature and salinity) can change when mixing with other water masses and when in contact with the surface, where heat and fresh water can be gained or lost. Here, we assume that large changes in water temperature and salinity (> 2 °C and 0.4 PSU over 20-km distance, respectively) measured around 5-m depth occurred when transiting between distinct water masses. For each cruise, water mixing events were manually annotated halfway along lines that connect two clusters of points in a temperature-salinity diagram. In Appendix A we illustrate a water mixing event.

### 2.2 Change-point analysis on point clouds

Now we present CytoSegmenter, our approach to change-point analysis on sequences of point clouds. Consider an ordered sequence of point clouds *x*_1_,…, *x_T_*. For every time index *t*, the point cloud 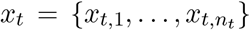 consists of *n_t_* points *x_t,i_* ∈ ℝ^*d*^. Assume that each point cloud *x_t_* consists of samples from an unknown probability distribution P_*t*_. We seek to estimate the times at which the distributions ℙ_*t*_ change in order to divide the sequence of point clouds into segments that are homogeneous in probability distribution. That is, if there are *m* change points, we seek to find the times *t*_0_:= 1 < *t*_1_ < *t*_2_ < · · · < *t_m_* < *t*_*m*+1_:= *T* + 1 for which ℙ_*t*_ ≠ ℙ_*t*+1_. We will do this by estimating the locations of a large number of potential change points and then using a penalty to estimate which set of possible change points is correct.

#### Change-point estimation

A natural way of segmenting a sequence of point clouds into *m* + 1 parts is to define a notion of a mean in the space of point clouds and then to optimize the segment boundaries so that, within a segment, the point clouds are as close as possible to the mean point cloud in that segment. One way to look at a point cloud is to see it an empirical probability measure and to smooth it using a nonparametric density estimate. Gretton et al. (2012) explains that a nonparametric density estimate, such as the ones considered by Anderson et al. (1994), can be summarized with a single quantity, called an empirical mean element, living in a Hilbert space.

The concept of a positive semi-definite (PSD) kernel is key to the notion of a mean element. Here a kernel is a function 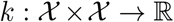 for some non-empty set 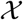 such that there exists a Hilbert space 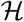 and a map 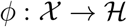 such that for all 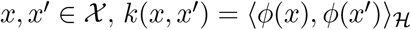 (Schölkopf and Smola, 2002). There are numerous benefits to the kernel viewpoint. For example, this approach can be applied to generic sets 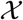, so long as a PSD kernel can be defined on 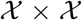. Numerous examples of PSD kernels can be found in the book of Shawe-Taylor and Cristianini (2004).

Equipped with this machinery, given a PSD kernel between two points in point clouds, we work with a point cloud via its corresponding mean element and leverage the Hilbert space structure to measure the distance between two points clouds via their corresponding mean elements. Then, applying that strategy another time, given a PSD kernel between two points clouds, we work with a series of point clouds via its corresponding mean element and leverage the Hilbert space structure to measure the distance of a point cloud to the mean element summarizing a collection of point clouds.

Concretely, let *k_x_*: ℝ^*d*^ × ℝ^*d*^ → ℝ be a kernel on points within each point cloud, with corresponding Hilbert space 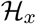 and feature map *ϕ_x_*. Given a point cloud *x_t_*, we define the empirical mean element as 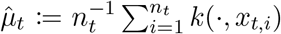. In addition, let 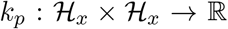 be a kernel on the empirical mean elements within each point cloud, with corresponding Hilbert space 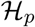 and feature map *ϕ_p_*. For a fixed number of change points *m* we obtain the change-point locations by minimizing a least-squares criterion in the Hilbert space 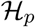:

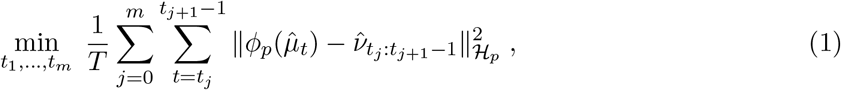

where 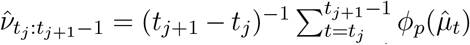 (Harchaoui and Cappé, 2007). We can evaluate the objective using only kernel evaluations (the “kernel trick”). The optimization can be done efficiently using a dynamic programming programming algorithm similar to the ones of Fisher (1958), Bellman (1961), and Kay (1993). We shall explain below how to set the number of change points *m*.

A straightforward implementation of the dynamic programming algorithm for a fixed *m* would make *O*(*T* ^2^) evaluations of the objective (1), which itself can be evaluated with a time complexity (max_*t*_ *n_t_*)^2^. To scale the approach to long series of data, we propose to use an approximation scheme, the Nyström method, to approximate a kernel (Williams and Seeger, 2000). This leads to a complexity in time that is linear in the number of points per point cloud rather than quadratic, allowing us to analyze series of tens or hundreds of millions of data points.

#### Parameter calibration and domain knowledge

We leverage auxiliary data with change-point annotations to set the number of change points in each sequence of point clouds. We assume here that we have (a) a parallel vectorial time series in which the number of change points is unknown but expected to be the same as in the time series of interest (in this paper, the physical data from the same cruise); and (b) a set of *n* similar vectorial time series for which the numbers of change points are known (in this paper, physical data from other cruises). The idea is that it can be easier to obtain change-point labels for time series of vectorial data than for time series of point cloud data.

We first use the *n* vectorial time series in (b) to calibrate the penalty parameter *α* from the penalty pen(*α, m*) = *α*(*m* + 1)(2 log(*T*/(*m* + 1)) + 5)*/T* in the penalized change-point problem of Lebarbier (2005). We set the value of *α* so that, across all *n* sequences, the difference between the true number of change points and the estimated number of change points in these vectorial sequences is minimum. Using this estimated *α*, we then run the method of Lebarbier (2005) on the corresponding parallel series of vectorial data in (a). This provides us with an estimate of the number of change points *m* in the corresponding sequence of point clouds, which we use when minimizing the objective in (1).

The other parameters that need to be calibrated are the parameters of the two kernels we use. We choose the kernel on datapoints *k_x_* to be the Gaussian radial basis function (RBF) kernel, which satisfies the assumptions of Sriperumbudur et al. (2008) for the theory of kernel embeddings of distributions to hold. As for the kernel *k_p_* on point clouds, we also use a Gaussian RBF kernel. For each RBF kernel we set the bandwidth to the median pairwise distance between inputs, a common rule of thumb. We perform the Nyström approximation with the projection of the kernel *k_x_* onto a subspace of size 128 (chosen as a reasonable balance between the accuracy of the kernel approximation and the runtime). We select the quadrature points by quantizing the data into a codebook of size 128 using 100 iterations of *k*-means.

In order to target phytoplankton community shifts associated with larger-scale oceanographic features, such as mesoscale eddies (~100 km) and gyre boundaries, and to avoid generating a large number of change points associated with high frequency variability, we set the minimum distance between change points to be 5 samples, which represents 15 min or roughly 5 km for a ship moving at 10 knots. In Appendix B we include the results of a sensitivity analysis of the parameters of our method. We find that the results are rather insensitive to the parameter values we set.

#### Code

The code for this paper was written in Python, builds upon Scikit-learn and Faiss (Pedregosa et al., 2011, Johnson et al., 2019) and may be found online at http://github.com/cjones6/cytosegmenter. All of the analysis, including the feature generation and change-point estimation, takes under 20 min total to run on a machine with an Intel i9-7960X processor, an Nvidia Titan Xp GPU, and 128GB of memory.

## 3 Results

In applying CytoSegmenter, the kernel change-point method described in Section 2.2, to SeaFlow data, we aim to answer the following questions:

1. Does the method successfully estimate changes in the distribution of phytoplankton communities?
2. Do the estimated change points in the biological data coincide with change points in the physical environment?
3. Can we leverage additional data on the physical environment from other cruises to estimate the number of change points?

For the first two questions we focus on a single cruise, KOK1606. As the number of change points in a cruise can be subjective, we initially fix the number of change points to ten. For the third question we expand the analysis to all cruises and estimate the number of change points in each cruise.

### Changes in distribution

We first aim to assess whether CytoSegmenter successfully estimates changes in the distribution of phytoplankton along the course of a cruise. To do so, we run the change-point estimation method on the biological measurements taken on the KOK1606 cruise when fixing the number of change points to 10. The locations of the resulting estimated change points are non-uniformly distributed across the cruise track (Fig. 2).

**Figure 2:**
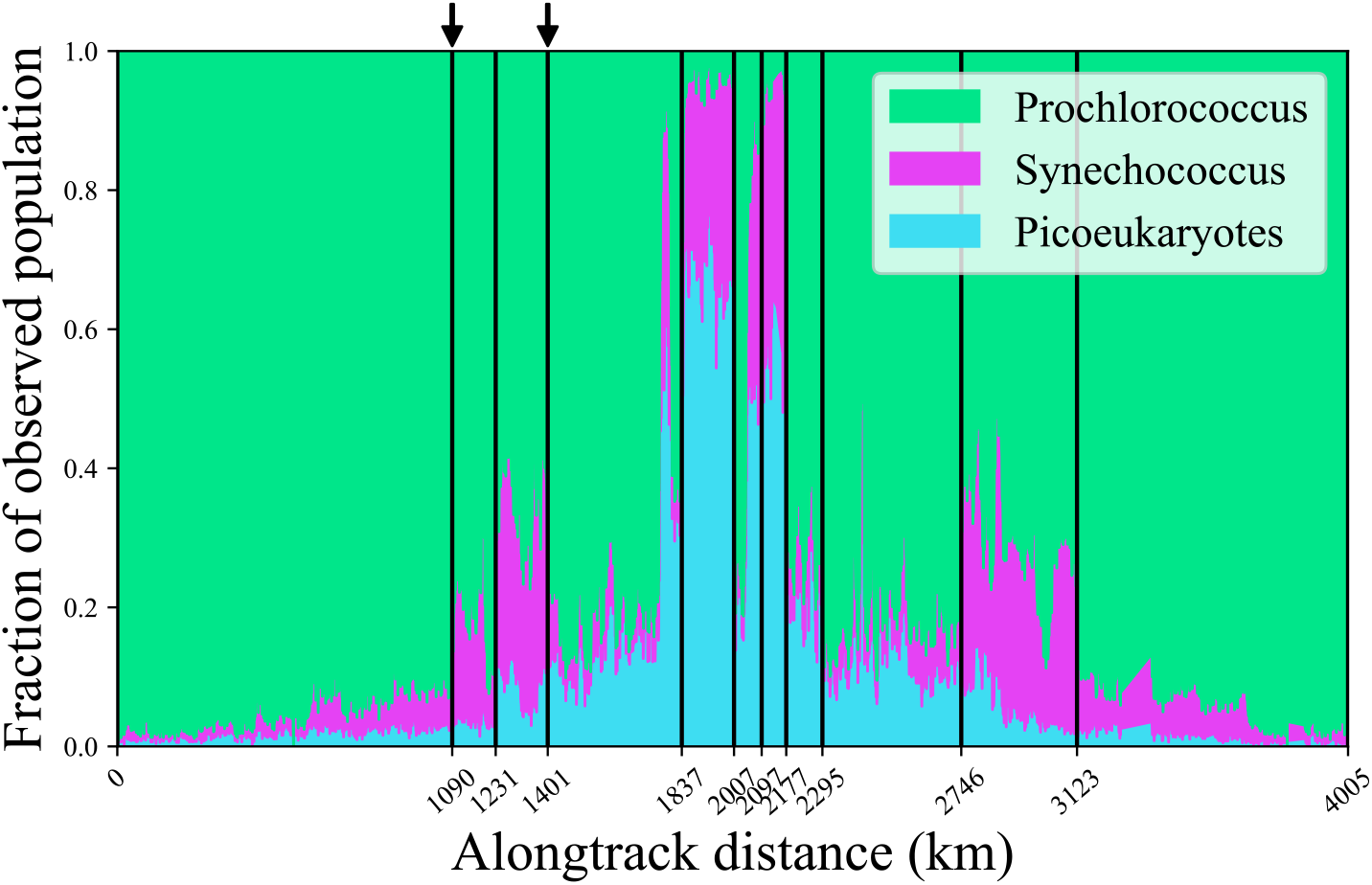
Estimated change points in the biological data from the KOK1606 cruise overlaid on the phytoplankton distribution observed during the cruise. The arrows at the top of the figure denote the change-point locations examined in Fig. 3.

We assess the quality of the estimated change points in three ways. First, Fig. 2 overlays the estimated change points on a plot of the phytoplankton distribution observed at each 3-minute time period throughout the course of the cruise. The plot illustrates that the algorithm locates large, abrupt changes in the phytoplankton community structure. For example, the algorithm detected the large increase from 5% *Synechococcus* to 18% *Synechococcus* at approximately 1090 km along the cruise track.

Next, for the change points at 1090 km and 1401 km we plot in Fig. 3a-3d the distribution of the log of either the chlorophyll or phycoerythrin measurements vs. the log of light scatter 5 km before and after the estimated change points. Recall that the cell labels were not available to our method; only the (raw) cell measurements were. From the plots we can see that there is a sudden increase between 1085 km and 1095 km in the fraction of cells that have light scatter between 10^4^ and 10^5^ and chlorophyll fluorescence between 10^3^ and 10^4^. From the labels, we can see that this corresponds to an increase in the fraction of *Synechococcus*. We can also see that from 1396 km to 1406 km there is a large drop in the fraction of cells that have light scatter between 10^4^ and 10^5^ and chlorophyll fluorescence between 10^4^ and 10^5^. This corresponds to a decrease in the fraction of *Synechococcus*. This decrease in *Synechococcus* is more pronounced when examining the distribution of the log of phycoerythrin vs. the log of light scatter (Fig. 3e-3f).

**Figure 3:**
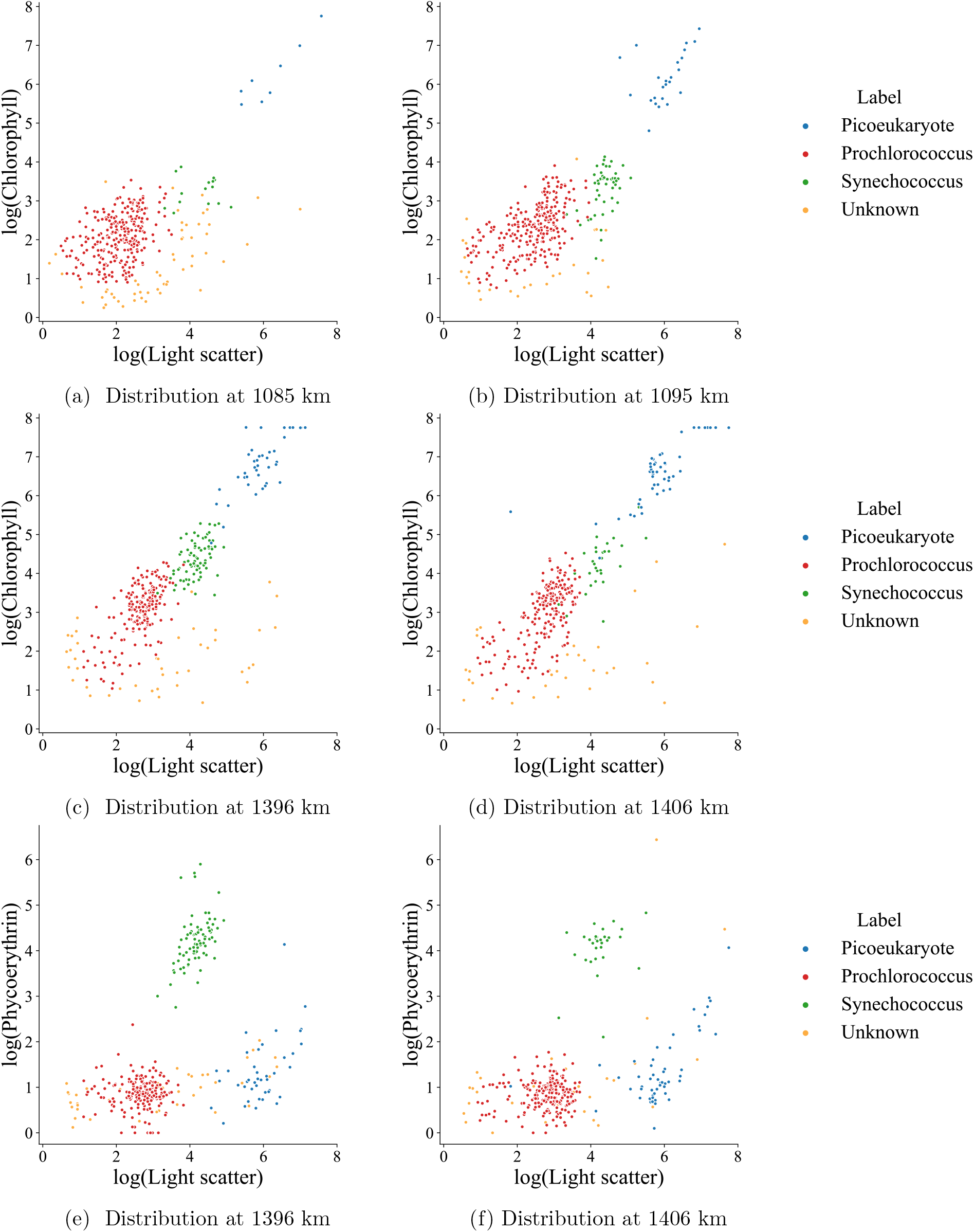
Distribution of the log of the chlorophyll or phycoerythrin measurements and the log of the light scatter measurements of phytoplankton five kilometers before and after estimated change points at 1090 km and 1401 km along the cruise track (KOK1606 in Table 1).

Finally, a distinctive feature of the KOK1606 cruise is that the northward and southward cruise trajectories were identical, occurring over a period of 3 weeks (see Fig. 4, which displays the latitude and longitude of the research vessel as a function of the alongtrack distance). This provided us with a method to check the efficacy and consistency of our method, as we expect that change points detected on northbound transect should be similarly detected on the southbound leg. The first change point, at 1090 km, is only 31 km in space from the last change point, at 3123 km. The fourth change point, at 1837 km, is 7 km from the fifth change point, at 2007 km. Moreover, the sixth and seventh change points, at 2097 km and 2177 km, are 24 km apart and the third and ninth change points, at 1401 km and 2746 km, are 59 km apart. Given that the ocean is a dynamic environment subject to constant movement driven by ocean currents, it is likely that the detected change points could shift in space. Surface mean current speeds in the region of the Pacific sampled by the KOK1606 cruise are ~ 0.1 m/s (Lumpkin and Johnson, 2013), which could drive a displacement of ~ 60 km over the course of a week. Spatial shifts in the ocean temperature during the cruise (cf. Fig. 5) support this hypothesis. The close spacing between our outbound and return change points confirms that our method is able to detect change points sampled more than once.

**Figure 4:**
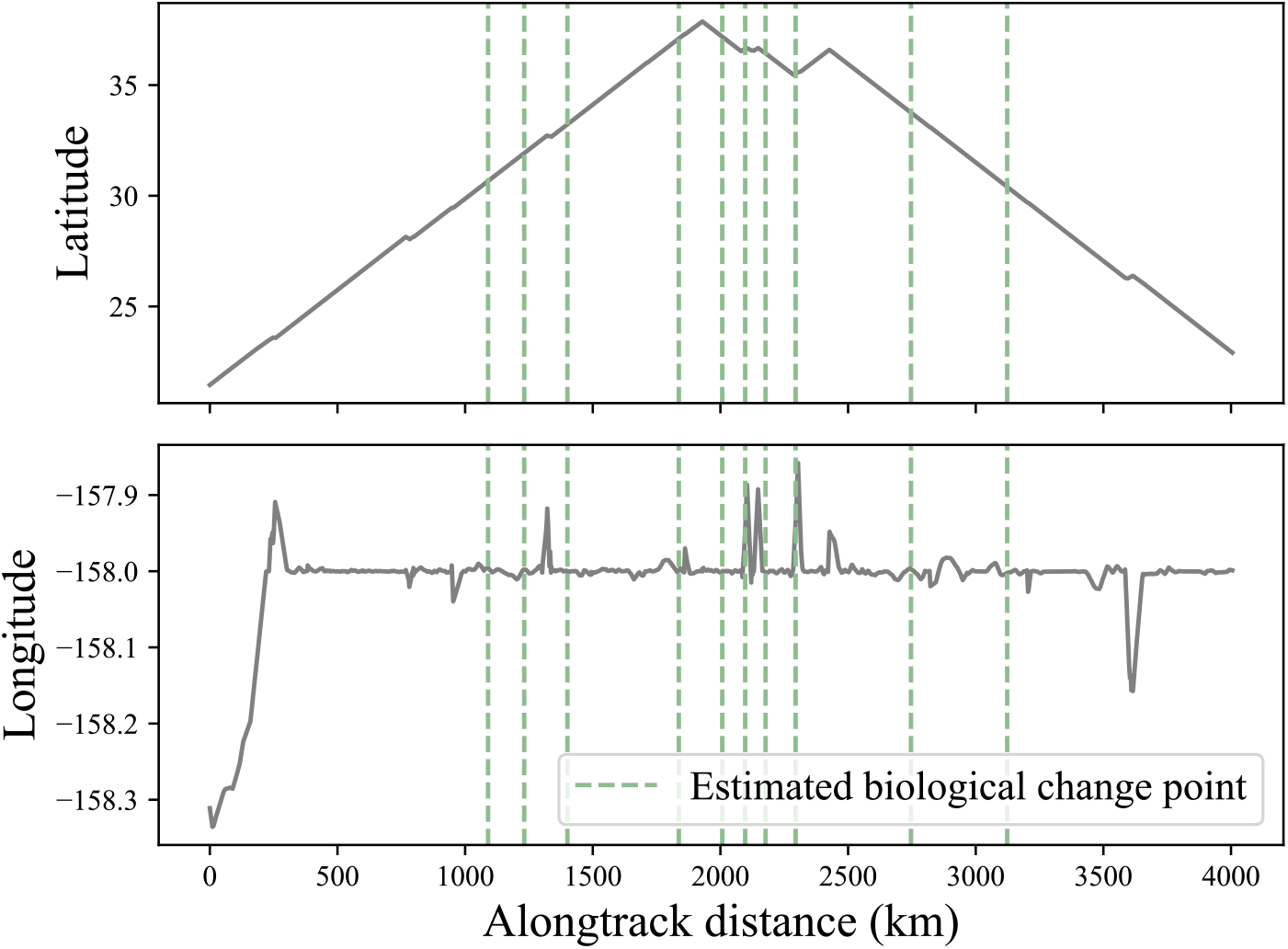
Estimated change points in the biological data from the KOK1606 cruise overlaid on the latitude and longitude of the ship over the course of the cruise.

**Figure 5:**
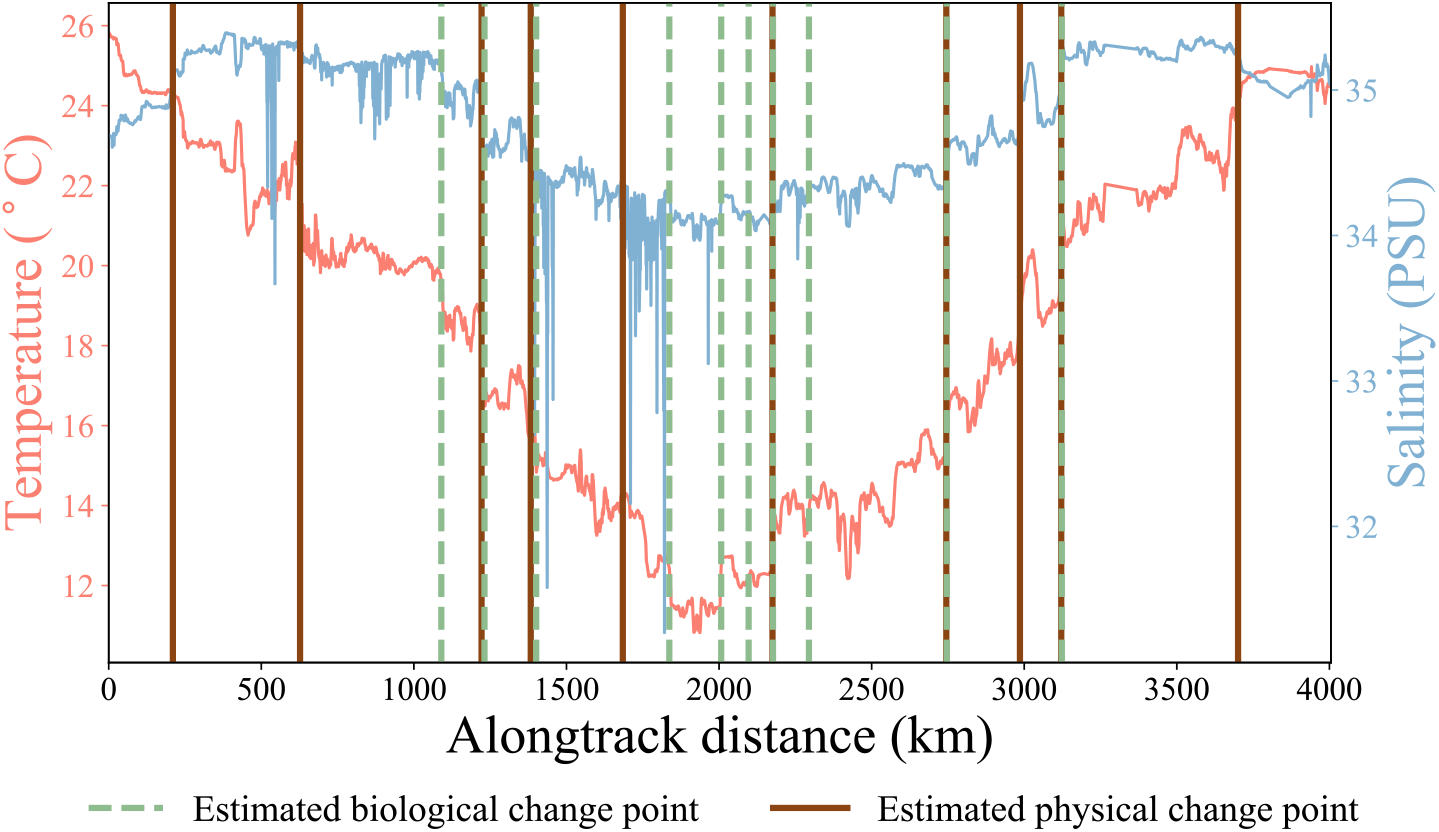
Estimated change points in the biological and physical data from the KOK1606 cruise overlaid on the temperature and salinity data recorded during the cruise.

### Correspondence between biological and physical change points

The distribution of individual phytoplankton species is a reflection of that species’ environmental niche (Hutchinson, 1957), defined as the range of environmental parameters within which a species can subsist. Niches can be controlled by physicochemical factors such as temperature and nutrient availability, as well as biotic processes such as competition and predation. In order to better understand the controls on phytoplankton distributions, we need to gain a better understanding of the balance between physical and biological controls in driving shifts in species’ distributions, which are reflected as shifts in overall phytoplankton community structure. By comparing the location of physical change points (based on surface temperature and salinity) and biological change points (based on SeaFlow data), we can begin to understand how important changes in the physical environment control phytoplankton community structure. When physical and biological change points coincide, that would suggest that physical processes are driving the observed community shift, whereas biological change points that do not coincide with physical change points likely reflect community shifts driven by biotic processes.

Fig. 5 plots the estimated change points overlaid on the temperature and salinity measurements throughout the cruise. The estimated physical and biological change points are within 20 km of each other 50% of the time. However, even when they do not quite coincide, the estimated biological change points are associated with large changes in temperature and salinity. These results suggest that shifts in phytoplankton community structure are largely associated with corresponding shifts in physical ocean properties. The results support previous work that showed that water masses play an important role in structuring phytoplankton communities (e.g., d’Ovidio et al., 2010).

### Estimation of the number of change points

Now we examine the results from estimating the number of change points. For each of the 12 cruises from Table 1 we estimate the parameter *α* based on the other 11 cruises, thereby obtaining one value of *α* per cruise. When considering the grid of *α*’s 0, 0.01, 0.02,…, 1 in the penalty of Lebarbier (2005), the value chosen is always either 0.12 or 0.13. The corresponding number of estimated physical change points for each cruise is presented in the left panel of Fig. 6. In this case the correlation between the number of estimated change points and the number of annotated change points is 0.63. Moreover, the number of estimated change points for 10 of the 12 cruises is within a factor of two of the number of annotated change points. This suggests that when annotated physical data for a given cruise is unavailable, the number of change points might be reasonably estimated from annotations of other cruises. In Appendix C we compare this approach to that of Harchaoui and Lévy-Leduc (2007) and demonstrate the superiority of this approach.

**Figure 6:**
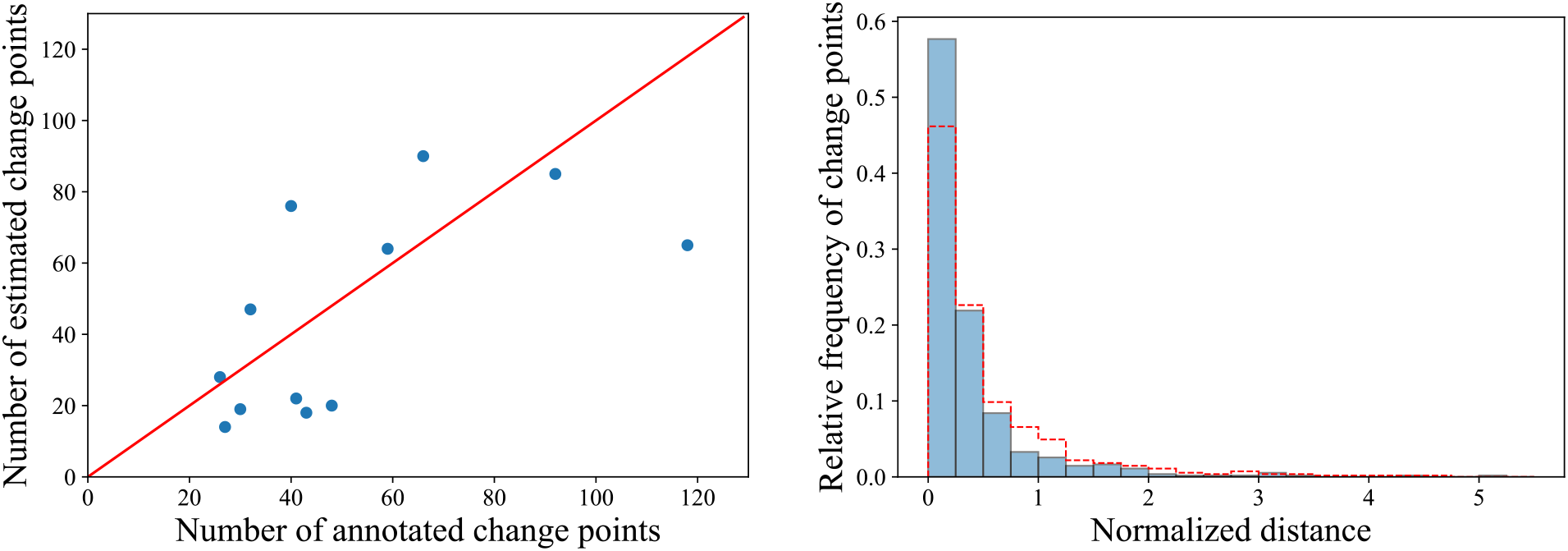
Estimated and annotated number of change points on the physical data (left) and histogram of the distances from each estimated biological change point to the nearest annotated physical change point for the same cruise, normalized by the average distance between annotated change points within the cruise (right). The diagonal red line in the plot on the left denotes the locations where the points would ideally lie. The red dashed line in the plot on the right indicates the histogram one would obtain from uniformly segmenting the cruises.

Using the estimated number of change points in the physical data from each cruise, we estimate the locations of change points in the biological data from each cruise. To assess the quality of the resulting estimated change points, we could compute the distance from each biological change point to the nearest annotated physical change point. That is, for a given cruise, if *t*_1_,…, *t_m_* and 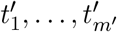 are the estimated and annotated change points, respectively, then in terms of distance traveled (by abuse of notation), we could compute for each estimated change point *t_j_* the value 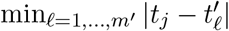. However, change points can be encountered at different rates during different cruises. Therefore, we normalize the aforementioned distances by the average distance between change points in each cruise. I.e., for each estimated change point *t_j_* we compute 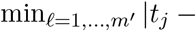 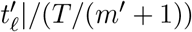, where *T* is the total distance traveled in the corresponding cruise.

The right panel of Fig. 6 plots a histogram of the normalized distances from each estimated biological change point to the nearest annotated physical change point. From the plot we can see that 58% of the estimated change points are within a normalized distance of 0.25 to the nearest annotated physical change point. In contrast, if we uniformly segmented each cruise this value would only be 46%. In fact, 22% of the estimated change points are within a normalized distance of 0.05 and 38% are within a normalized distance of 0.1, compared to 9% and 19% based on a uniform segmentation.

Finally, Fig. 7 displays the estimated and annotated change points for the KOK1606 cruise. The number of estimated change points is 64, whereas the number of annotated change points is 60.

**Figure 7:**
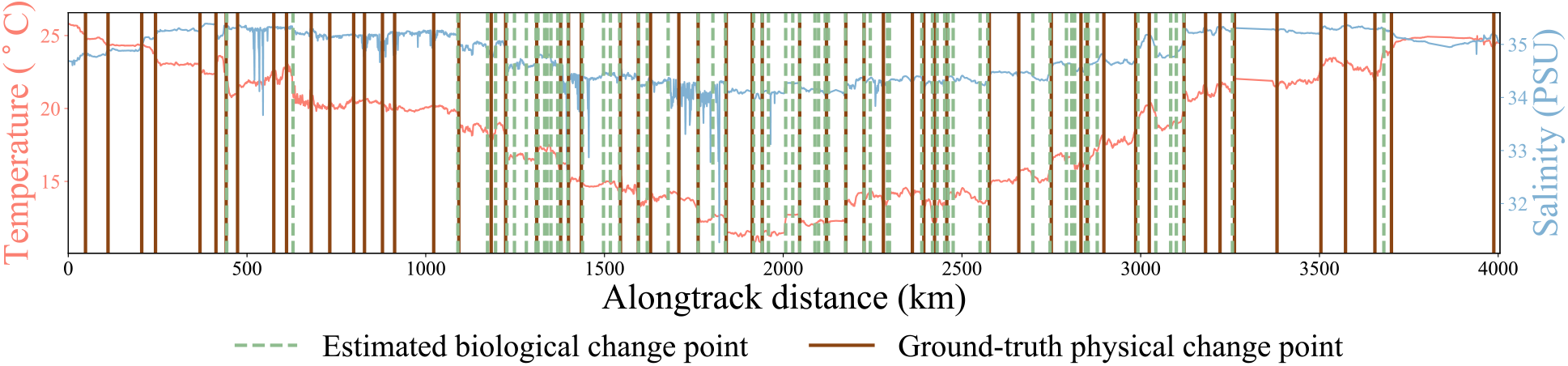
Estimated change points in the biological data and annotated change points in the physical data from the KOK1606 cruise overlaid on the temperature and salinity data recorded throughout the cruise.

## 4 Discussion

In this work we presented a kernel-based change-point detection method that applies to point cloud data. We applied the method to data collected by a shipboard flow cytometer during research cruises. The results suggest that the method is able to locate meaningful changes in the phytoplankton distribution. Furthermore, we found that changes in the distribution of phytoplankton generally co-occur with changes in temperature and salinity. This paved the way for estimating the number of change points in the biological data based on the number of change points in the physical data.

### 4.1 Comparison to other methods

A previously-proposed approach to the detection of changes in flow cytometry data is that of Hyrkas et al. (2015). This approach directly averages the points within each point cloud to obtain a single vector and then applies the multiple change-point detection method of Matteson and James (2014). The pre-processing step results in a significant loss of information. In particular, the higher-order moment information of each point cloud distribution is lost in the process. The CytoSegmenter approach we propose here is based on a nonparametric density estimate, which captures all moments of a point cloud distribution, hence capturing richer statistical information than the direct approach of Hyrkas et al. (2015).

An alternative empirical approach specific to flow cytometry data is based on the so-called cytometric diversity; see the paper of Li (1997) for an overview. Each point cloud is summarized into ataxonomical categories based on the bio-optical properties measured at the single cell level by flow cytometry. Cytometric diversity then expresses the organization and structure of a community, its richness in physiological and genetic variations. The Hill diversity measure of order *α* is defined as 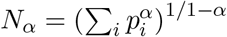, where *α* is usually 0, 1 or 2, and *p_i_* is the proportional abundance of the *i*-th category of phytoplankton. The maximum number of distinct categories depends on the resolution at which each optical property is measured. In the case of SeaFlow, light scatter, red and orange fluorescence are resolved to 2^16^ discrete values, which represent a maximum of 2^48^ categories for each point cloud. Changes in phytoplankton structure are then detected by computing the first or second derivative of the cytometric diversity indices, after smoothing out high frequency or random noise in the signal using a low-pass filter set by the user (Ribalet et al., 2010). While capturing the point cloud organization in a rather sophisticated way, this approach relies on ad hoc choices for the number of bins prior to the analysis, making further comparisons across datasets rather difficult.

### 4.2 Broader applicability of the method

We have developed a statistical method for identifying changes in the underlying structure of a sequence of multi-dimensional point clouds. The method can be applied to sequences collected over time and/or space, and is applicable even when the number of points per point cloud varies. Point cloud data also arises in other areas such as computer graphics and computer vision (see, e.g., Suard et al., 2005, Lézoray and Grady, 2012, Stumm et al., 2016). Here we have applied this method to identify shifts in phytoplankton community structure from a large dataset of underway flow cytometry measurements. We envision that this statistical framework will be widely applicable to datasets structured in a similar way. A range of broadly similar continuously sampling flow cytometers are currently commercially available (CytoSense, Imaging FlowCytobot, FlowCam) which collect cytometric data (e.g., scatter and fluorescence) as well as images of plankton cells from environmental samples. The images collected by these instruments are used to identify plankton species and to train machine learning models for automated classification (Sosik and Olson, 2007). The change-point detection method we have developed could be applied to such data in order to identify broad planktonic community shifts before fully identifying the images.

The approach we introduced in this paper is both fast and scalable. In future work it would be interesting to explore related online methods for change-point detection on point cloud data that could be used in real time during research cruises to help scientists develop an adaptive sampling approach.

## 5 Acknowledgements

We thank Chris Berthiaume and Dr. Annette Hynes for their help in processing and curating SeaFlow data. This work was supported by grants from the Simons Foundation (Award ID 574495 to F.R., Award IDs 329108, 426570SP and 549894 to E.V.A.). C.J. acknowledges support from NSF grant DMS-1810975. S.C. acknowledges support from a Moore/Sloan Data Science and Washington Research Foundation Innovation in Data Science Postdoctoral Fellowship. Z.H. acknowledges support from the CIFAR LMB program and NSF grant DMS-1810975. The authors are grateful to the eScience Institute at the University of Washington for supporting this collaboration.

## 6 Data Accessibility and Reproducibility

The physical and biological data may be found at https://doi.org/10.5281/zenodo.4289399, while the code is located at http://github.com/cjones6/cytosegmenter.

## 7 Author contribution statement

CJ, ZH, and SC conceived the ideas and designed the methodology; FR and EVA collected and curated the data; CJ analysed the data with the help of FR and SC; CJ led the writing of the manuscript. All authors contributed critically to the drafts and gave final approval for publication.

## A Water Mixing Events

We produced manual change point annotations in the physical data by locating water mixing events. Figure 8 displays an example of a water mixing event. We can see from the panel on the left that the temperature and salinity drop from 18.5 − 19°C and 35 − 35.1 PSU, respectively, to around 16.5 °C and 34.5 PSU over a distance of 15 km. We label the approximate midpoint of this drop as a change point. The mixing event can also be seen via a plot of the temperature vs. salinity (right panel of Fig. 8). In this plot the colors progress from red to violet as the distance along the cruise track increases. We can see that there are two clusters of temperature-salinity measurements, and the ship progressed from the upper-right cluster of points around 19°C and 35.05 PSU to the cluster of points around 16.7°C and 34.6 PSU. The change point we annotate (with the black X) is approximately halfway between the two clusters.

**Figure 8:**
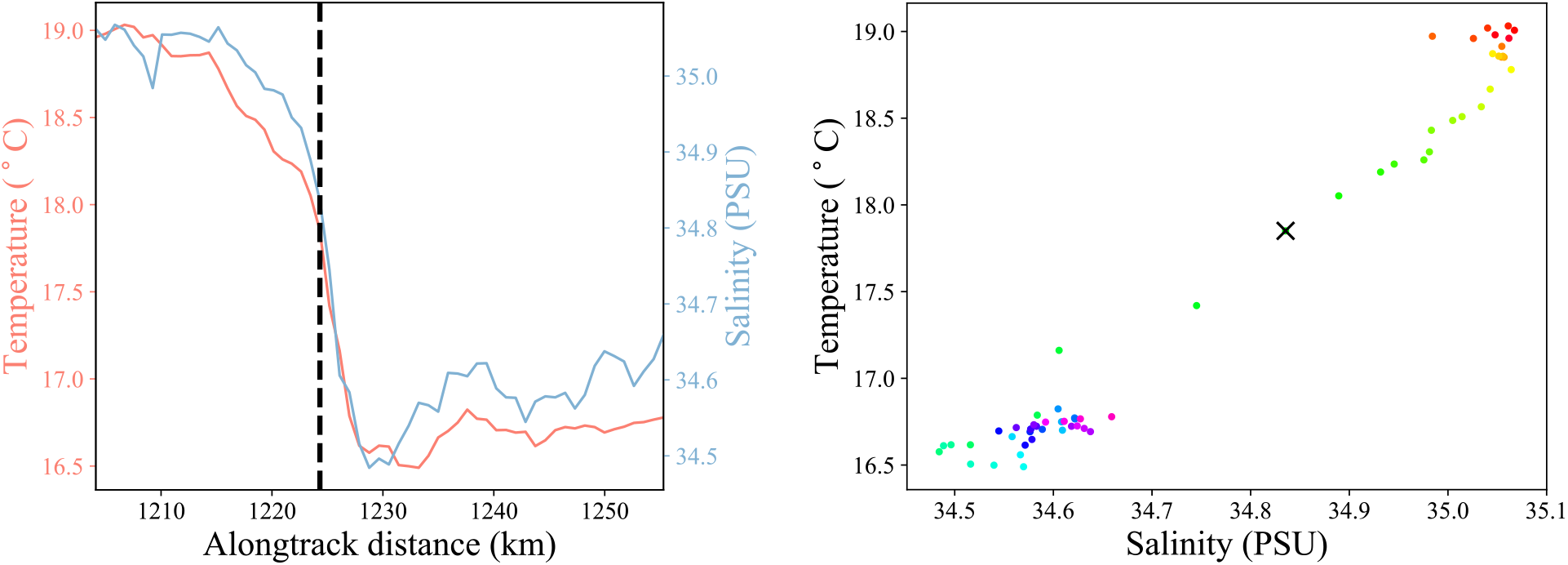
Annotation of a mixing between different water masses encountered during the KOK1606 cruise. The dashed black vertical line (left plot) and black X (right plot) denote the location of an annotated change point. The colors of the points in the plot on the right advance from red to violet as the distance traveled increases.

## B Sensitivity Analysis

We examined the sensitivity of the estimated change points to the parameter settings. We examined this for data from the KOK1606 cruise with 10 change points. The results are shown in Figure 9. In each plot we display in the background the distribution of phytoplankton at each point during the cruise (the same distributions as in Figure 2). In the foreground we display rows of colored boxes, denoting the locations of change points along the x-axis for a given parameter setting.^2^

**Figure 9:**
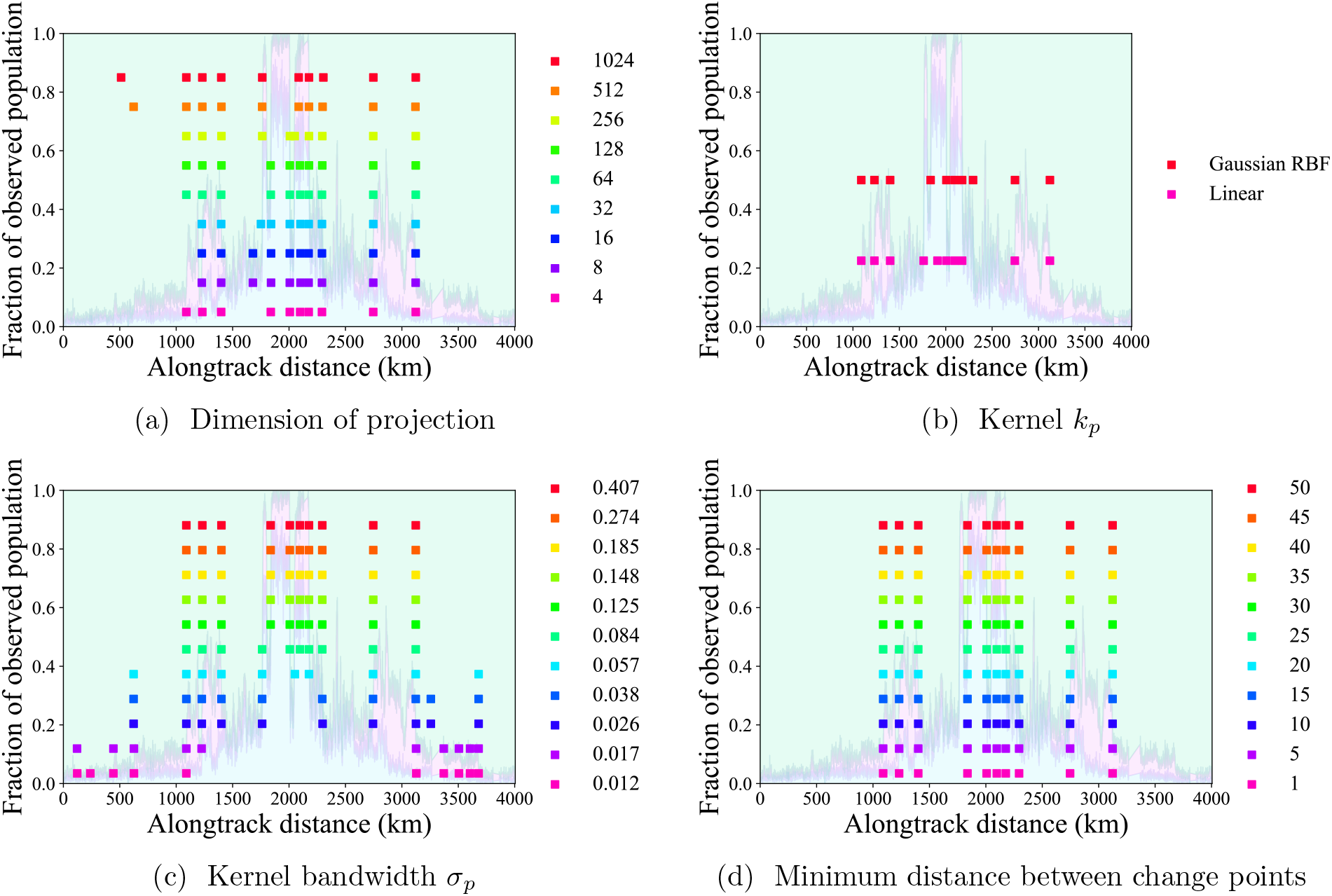
Sensitivity of the estimated change points in the biological data from the KOK1606 cruise to changes in the parameters of the method.

We first varied in Figure 9a the dimension of the projection onto the subspace. We find that when using projections with dimensions 4 through 128, nine of the ten change points lie within 4 km of each other. However, they do not agree on whether there is a change point at 1090 km, 1680 km, or 1751 km. With a projection of dimension 256 the method estimates there to be a change point at 1763 km rather than 1751 km and 2051 km rather than 2100 km. Despite these differences, we can see that there are large changes in distribution at each of these locations. Once the dimension is increased to 512 or 1024 the method begins to detect a smaller change in distribution closer to the beginning of the cruise.

Next we examined the effect of the kernel *k_p_*. Comparing the Gaussian RBF kernel to the linear kernel, we see from Figure 9b that the estimates differ by two change points. While the Gaussian RBF kernel puts change points at 1837 km and 2295 km, the linear kernel puts change points at 1762 km and 1911 km. Visually, there are large changes in the distribution of phytoplankton at all of these locations. The remainder of the estimated change points from the Gaussian RBF kernel are all within 1 km of the corresponding change points estimated with the linear kernel.

For the Gaussian RBF kernel we used the median pairwise distance rule-of-thumb by default, which resulted in a bandwidth of 0.148. In Figure 9c, we vary the quantity 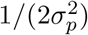 on a log scale, where the endpoints are determined by setting *σ_p_* to the 1st and 99th percentiles of the distances between inputs. For relatively small bandwidths (less than 0.084) the algorithm tends to pick up smaller changes in distribution closer to the start and/or end of the cruise. From 0.084 up through 0.407 the change point estimates do not differ by more than 1 km, with the exception of the fourth change point. With a bandwidth of 0.084 the fourth change point is estimated to be at 1762 km, whereas for the other bandwidths it is estimated to be at 1837 km.

Finally, we examined the effect of setting a minimum distance between change points in Figure 9d. The change points estimated with each minimum distance are the same.

## C An Alternative Approach to Estimating the Number of Change Points

An alternative approach to estimating the number of change points in a sequence is to use the rule of thumb of Harchaoui and Lévy-Leduc (2007). The rule of thumb says to choose the minimum number of change points *m* such that the ratio of successive objective values with *m* + 1 change points and *m* change points exceeds 1 − *ν* for some value *ν*. In our analysis the tuning we perform in this case is over the threshold for the ratio of successive objective values 1 − *ν* = 0.9, 0.901, 0.902,…, 0.99, and the value chosen is always 0.98. Figure 10 displays the resulting estimated vs. annotated number of change points in the physical data (cf. Figure 6). The correlation between the estimated and annotated number of change points is −0.14. In contrast, with the penalty-based approach described in Section 2.2 the correlation was 0.63. These results suggest that the method of Lebarbier (2005) is promising in this context while the rule of thumb of Harchaoui and Lévy-Leduc (2007) is ineffective.

**Figure 10:**
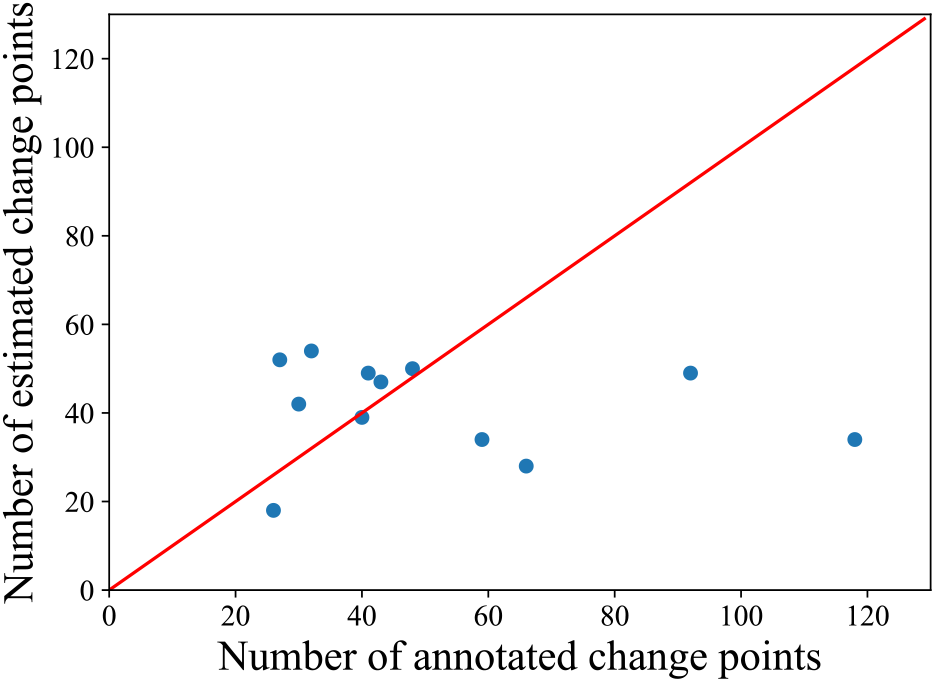
Estimated and annotated number of change points in the physical data based on the rule of thumb from Harchaoui and Lévy-Leduc (2007). The diagonal red line denotes the locations where the points would ideally lie.

## Footnotes

1 NOAA OI SST V2 data provided by the NOAA/OAR/ESRL PSL, Boulder, Colorado, USA, from their website at https://psl.noaa.gov/.

2 Note that the vertical locations of the boxes do not have meaning, aside from corresponding to different parameter settings.

